# Glioblastoma is associated with extensive accelerated brain ageing

**DOI:** 10.1101/2022.08.24.505089

**Authors:** Anna P. Ainslie, Myrthe Klaver, Daniëlle C. Voshart, Emma Gerrits, Bart J.L. Eggen, Steven Bergink, Lara Barazzuol

## Abstract

Progressive neurocognitive dysfunction is the leading cause of a reduced quality of life in patients with primary brain tumours. Understanding the mechanisms underlying cognitive impairments that occur in response to a brain tumour and its treatment is essential to improve patients’ quality of life. Here, we show that normal-appearing non-tumour brain regions of patients with glioblastoma display hallmarks of accelerated ageing and share multiple features with Alzheimer’s disease patients. Integrated transcriptomic and tissue analysis shows that normal-appearing brain tissue from glioblastoma patients has a significant overlap with brain tissue from Alzheimer’s disease patients, revealing shared mitochondrial and neuronal dysfunction, and proteostasis deregulation. Overall, the brain of glioblastoma patients undergoes Alzheimer’s disease-like accelerated ageing, providing novel or repurposed therapeutic targets for managing brain cancer-related side effects.

## Introduction

Primary brain tumours account for over 20% of paediatric, and 1.4% of adult cancers^1,2^. Recent advances in brain tumour treatment have led to an increase in the proportion of long-term survivors depending on tumour subtypes and paediatric versus adult patients. In adult cases the median overall survival ranges between 14.6 months for grade IV glioblastoma (GBM) to 13.8 years for lower grade II glioma^3^, while in the paediatric population the 5-year survival rate has increased to over 75%^4^. Nearly all patients with primary brain tumours develop debilitating neurocognitive dysfunction, resulting in a reduced quality of life, and educational and occupational attainment^5,6^. Importantly, this impairment in neurocognitive function is irreversible and progressive in nature, in some cases developing long after completion of treatment^6,7^. It affects various neurocognitive domains, in particular memory, processing speed and executive function, resulting in dementia in 5% of cases^8,9^.

Many factors play a role in the development of neurocognitive dysfunction, including tumour type, size and location, type of cancer treatment, which often involves a combination of surgery, radiotherapy and chemotherapy, as well as individual genetic variation^3^. For instance, genome-wide association studies have reported a number of single nucleotide polymorphisms (SNPs) that are associated with worse neurocognitive outcome in patients with adult brain tumours after treatment with chemotherapy and radiotherapy^3,10^. Interestingly, several of these SNPs are in genes implicated in Alzheimer’s disease (AD), such as apolipoprotein E (*ApoE*), catechol-O-methyltransferase (*COMT*) and brain-derived neurotrophic factor (*BDNF*)^3^.

Despite the major impact of neurocognitive dysfunction, the mechanisms mediating this cognitive decline in patients with brain tumours remain largely not understood. Here, we investigated how the human brain responds to cancer and its treatment by performing comparative transcriptional profiling of post-mortem brain samples from patients with GBM. We found that normal-appearing non-tumour brain regions from GBM patients display extensive mis-regulation of genes involved in mitochondrial and neuronal function. Gene set enrichment analyses and a direct comparative transcriptomic analysis with an independent cohort of brain samples from AD patients revealed a significant overlap with AD. Furthermore, histological and protein analyses showed an increase in hallmarks of ageing and loss of proteostasis in normal-appearing brain regions from GBM patients. Overall, these data indicate that the brain of GBM patients undergoes accelerated ageing with a similar biological trajectory to AD.

## Results

### Normal-appearing brain tissues of GBM patients display major transcriptomic alterations

To identify the consequences of a brain tumour and its treatment on the healthy brain, we performed a comparative transcriptomic analysis of human post-mortem brain samples derived from healthy subjects and patients with GBM treated with chemotherapy and radiotherapy. We obtained non-tumour normal-appearing brain tissue from five GBM patients (NA-GBM), and age-matched and region-matched brain tissue from five unaffected control individuals (Figure 1a, Supplementary Figure 1a). The NA-GBM patient samples were collected from regions distant from the tumour to minimise the influence of the tumour (Supplementary Figure 1a, Table 1). From these samples, RNA was isolated and bulk RNA-sequencing was performed. 917 upregulated differentially expressed genes (DEGs) and 701 downregulated DEGs were detected when comparing the NA-GBM to control samples (Supplementary Data 1, Supplementary Figure 1b,c). Cell deconvolution analysis^11,12^ revealed a signature that indicated a high proportion of excitatory neurons across all samples (Figure 1b). Importantly, the astrocyte-like signature, characteristic of GBM cells^13^, was similarly low in all the samples suggesting minimal to no infiltration of cancer cells in our samples (Figure 1b). Immunostaining for Nestin, a protein strongly associated with GBM^14^, showed similar levels across all samples, confirming the absence of detectable tumour material in the samples (Figure 1c,d).

**Figure 1.**
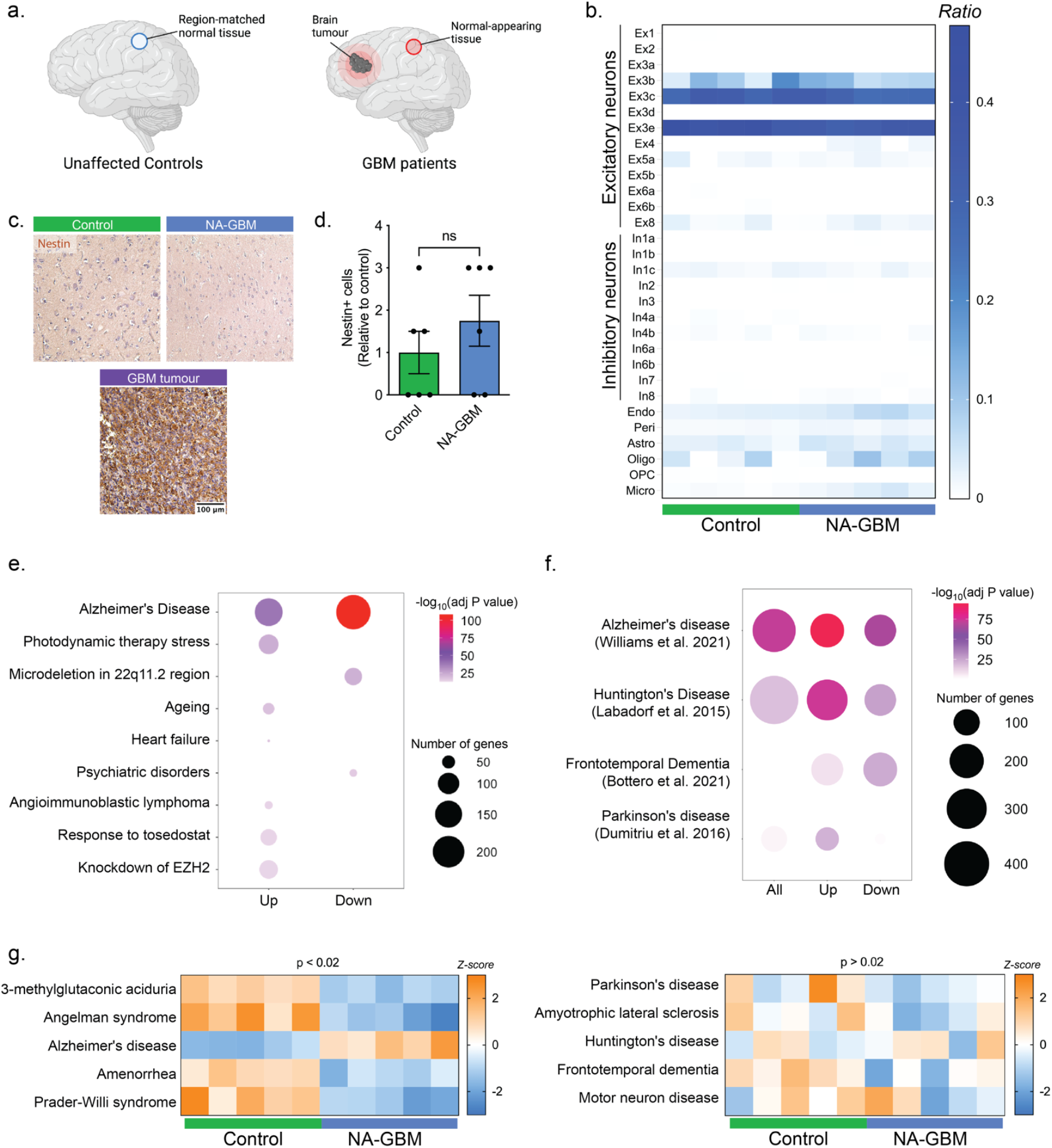
Transcriptomic analysis of normal-appearing GBM patient brain tissue. a. Non-tumour normal-appearing GBM (NA-GBM) patient brain tissue samples were analysed and compared to region-matched healthy tissue from unaffected control individuals. b. Cell deconvolution analysis of the unaffected control (n = 5) and NA-GBM (n = 5) data sets using the CIBERSORTx analytical tool^11^. Heat map indicates inferred ratio of cell types. Endo = endothelial, Peri = pericytes, Astro = astrocytes, Oligo = oligodendrocytes, OPC = oligodendrocyte precursor cells, Micro = microglia. c. Immunohistochemistry staining for Nestin in unaffected control, NA-GBM and GBM tumour samples. Scale bar = 100 µm. d. Quantification of Nestin staining, showing number of Nestin+ cells per 9 mm^2^ in unaffected control (n = 6) and NA-GBM samples (n = 6), relative to control (p = 0.5238, Mann-Whitney U). Data are represented as mean ± SEM. e. Results of enrichment analysis, comparing the NA-GBM vs unaffected control DEG list to the transcriptomic data from the “Chemical and genetic perturbations” data set from the Molecular Signatures Database (MSigDB). The top hits are plotted for the upregulated DEGs and the downregulated DEGs (-log_10_(adjusted P value) > 12). Scale bar indicates -log_10_(adjusted P value) and dot size represents number of overlapping genes. f. Manual comparison of GBM vs unaffected control DEG list to DEGs from selected neurodegenerative disease transcriptomic studies^16-19^. Scale bar indicates -log_10_(adjusted P value) and dot size represents number of overlapping genes. g. PGSEA analysis for unaffected control (n = 5) and NA-GBM (n = 5) samples against the Jensen disease database^41^, scale bar indicates the Z-score for each sample. Left plot shows the top 5 hits (p < 0.02), and the right plot shows the results for selected neurodegenerative diseases (p > 0.02).

**Table 1.** Description of post-mortem brain tissue samples used for bulk RNA-sequencing.

| Sample type | Sample label | Age | Sex | Brain region | PMI | Tumour location | Colour | Radiotherapy | Chemotherapy | Lipofuscin and IHC | WB |
| --- | --- | --- | --- | --- | --- | --- | --- | --- | --- | --- | --- |
| Unaffected control individuals | Control_1 | 54 | Male | Right Dorsolateral Prefrontal cortex | 9 |  |  |  |  | Y | Y |
|  | Control_2 | 54 | Female | Left Anterior Cingulate | 6 |  |  |  |  | Y | Y |
|  | Control_3 | 50 | Female | Left Dorsolateral Prefrontal cortex | 15 |  |  |  |  | N | N |
|  | Control_4 | 49 | Female | Left Posterior Cingulate | 26 |  |  |  |  | N | Y |
|  | Control_5 | 60 | Male | Left Dorsolateral Prefrontal cortex | 37 |  |  |  |  | Y | Y |
| GBM patients | NA_GBM_1 | 55 | Female | Left Anterior Cingulate | 21 | Left hemisphere | Purple | Y | Y | Y | N |
|  | NA_GBM_2 | 60 | Female | Left Dorsolateral Prefrontal cortex | 21 | Right frontal lobe | Yellow | Not available | Not available | N | Y |
|  | NA_GBM_3 | 57 | Female | Left Dorsolateral Prefrontal cortex | 8 | Pons and medial, left cerebellar white matter | Blue | Y | Y | N | Y |
|  | NA_GBM_4 | 49 | Female | Left Posterior Cingulate | 3 | Right posterior parietal | Red | Y | Y | Y | Y |
|  | NA_GBM_5 | 62 | Male | Left Dorsolateral Prefrontal cortex | 7 | Left posterior superior temporal lobe | Green | Y | Y | N | Y |
| AD patients | AD_1 | 44 | Male | Left Dorsolateral Prefrontal cortex | 14 |  |  |  |  | N | Y |
|  | AD_2 | 60 | Female | Left Dorsolateral Prefrontal cortex | 15 |  |  |  |  | N | Y |
|  | AD_3 | 59 | Female | Left Dorsolateral Prefrontal cortex | 19 |  |  |  |  | N | N |
|  | AD_4 | 56 | Male | Left Dorsolateral Prefrontal cortex | 4 |  |  |  |  | Y | Y |
|  | AD_5 | 59 | Male | Right Dorsolateral Prefrontal cortex | 13 |  |  |  |  | N | N |

Gene ontology (GO) analysis showed many significantly enriched terms in the upregulated gene clusters. The most significantly enriched terms were those involved in inflammation, such as a positive regulation of cytokine production (Supplementary Figure 1d, Supplementary Figure 2a). The most significantly enriched terms in the downregulated gene clusters were involved in oxidative phosphorylation, cellular respiration, and proton transmembrane transport (Supplementary Figure 1d, Supplementary Figure 2a). Clusters 1 and 6 also contained an enrichment of downregulated genes involved in neuron projection development, as well as regulation of long-term neuronal synaptic plasticity (Supplementary Figure 1d). Overall, the GO analysis showed an upregulation of genes involved in inflammation, and a downregulation of genes involved in oxidative phosphorylation and neuronal development.

Next, we asked whether the gene expression changes identified in NA-GBM brain tissue display an overlap with other disease conditions and compared our DEG dataset with previously published datasets, using enrichment analysis against the Molecular Signatures Database. This study showed a significant number of overlapping genes with the Blalock *et al*. Alzheimer’s disease (AD) transcriptomic study (Figure 1e)^15^. The top hit for both the downregulated and the upregulated genes was a publication studying the gene expression in AD patients^15^, with a statistically significant overlap of 158 upregulated genes (p = 2.50E-39) and an overlap of 234 downregulated genes (p = 5,27E-109) (Figure 1e). Manual comparison between the NA-GBM DEGs and selected transcriptomes from multiple neurodegenerative diseases^16-19^ showed that AD had the most significant overlap of both up- and down-regulated genes as well (Figure 1f). Additionally, parametric gene set enrichment analysis (PGSEA) comparing the NA-GBM DEGs against the Jensen disease database showed that genes associated with Alzheimer’s disease positively and significantly correlate with the DEGs we found in NA-GBM tissue (Figure 1g). Conversely, genes associated with other neurodegenerative diseases (NDs) do not correlate significantly with the DEGs found in NA-GBM tissue (Figure 1g).

### Identification of shared transcriptomic features between normal-appearing GBM brain tissue and Alzheimer’s disease

To test the overlap in gene expression between NA-GBM tissue and AD directly, five additional AD patients were included in our study for differential gene expression analysis. Cell deconvolution analysis confirmed a similarly high proportion of excitatory neurons in the AD samples (Supplementary Figure 3a). 531 DEGs were upregulated and 631 were downregulated when comparing AD patient samples to unaffected control individual samples (Supplementary Data 1). By contrast, only a total of 59 DEGs were differentially regulated when comparing AD and NA-GBM, suggesting similarities between the two datasets (Supplementary Figure 3b). Comparing the DEGs in the NA-GBM samples versus control samples and the AD samples versus control samples showed an overlap of 615 DEGs, of which 272 upregulated and 343 downregulated genes (Figure 2a, Supplementary Data 1). Overall, there is a significant overlap between the DEGs in NA-GBM patient samples and the DEGs in AD patient samples (Figure 2b). Unbiased Euclidean clustering confirmed this as the control samples clearly segregated while the AD and NA-GBM samples were nearly indistinguishable (Figure 2c), in line with the PCA analysis (Supplementary Figure 3c). Additionally, the PCA plot revealed that sample clustering is not correlated with age (Supplementary Figure 3d). Together, our findings indicate that the gene expression profile of non-tumour tissue in GBM patients closely resembles AD.

**Figure 2.**
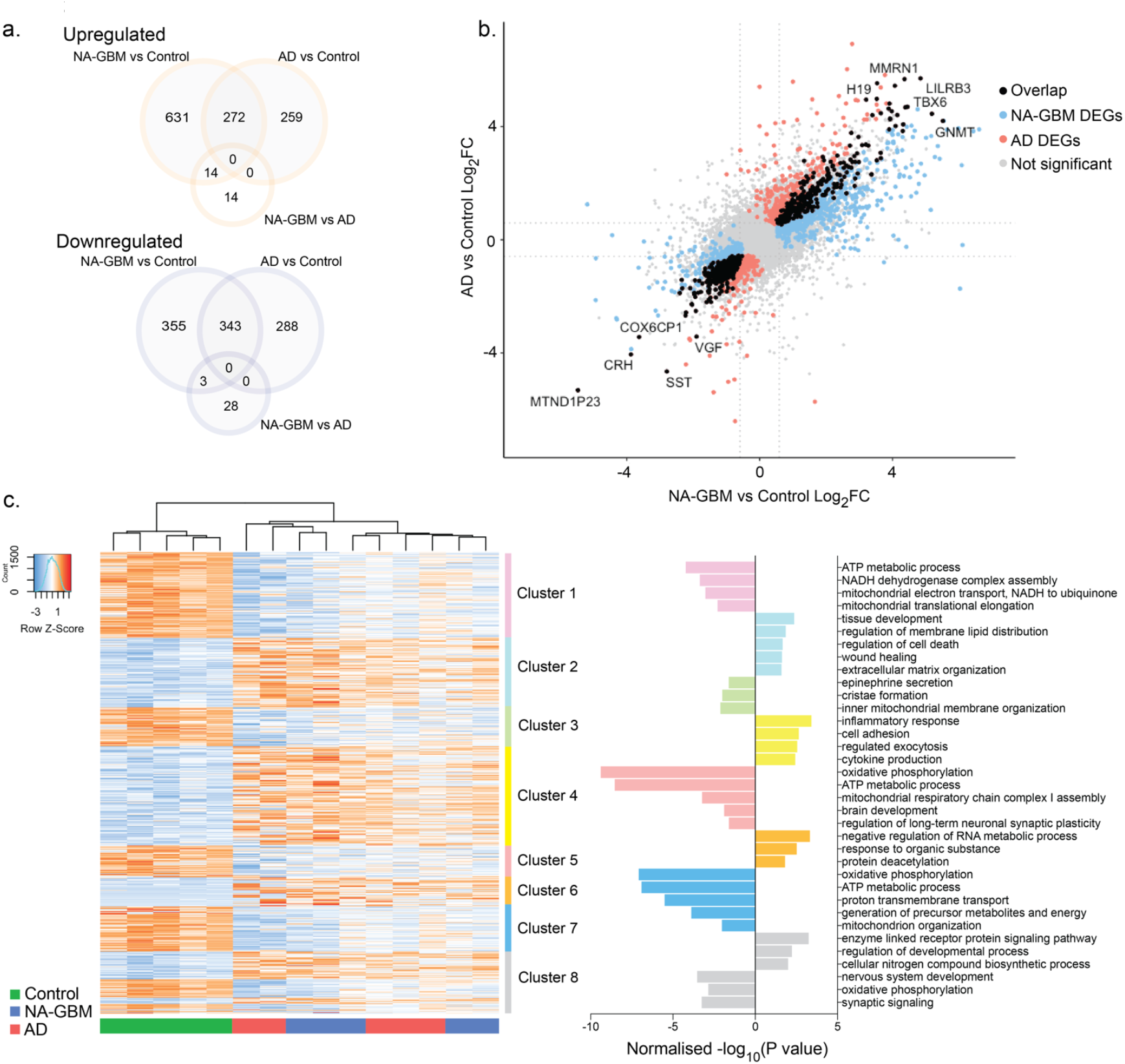
Overlapping gene expression patterns in normal-appearing GBM and AD patient brain tissue. a. Venn diagram of overlapping upregulated and downregulated DEGs between NA-GBM vs unaffected control, AD vs unaffected control and NA-GBM vs AD data sets, determined on the basis of fold change > 1.5 and FDR < 0.05. b. Four-way plot showing 615 overlapping DEGs between NA-GBM vs unaffected control and AD vs unaffected control data sets, determined on the basis of fold change > 1.5 and FDR < 0.05. c. Clustered heat map of DEGs when comparing NA-GBM patient samples and AD patient samples together to unaffected controls (fold change > 1.5 and FDR < 0.05). GO analysis (Biological Processes) of individual clusters using g:Profiler, normalised -log_10_(P value) is plotted on the right.

To evaluate the extent of transcriptomic similarity, the NA-GBM and AD data sets were combined and compared with the control samples for differential gene expression analysis. We detected 976 upregulated DEGs and 954 downregulated DEGs (Supplementary Data 1). GO analysis showed many significantly enriched terms in the upregulated gene clusters involved in different biological processes, the top terms include inflammation, regulation of lipid distribution, regulation of cell death, and extracellular matrix organisation (Figure 2c). Many significantly enriched terms in the downregulated gene clusters are involved in mitochondrial membrane organisation, oxidative phosphorylation and nervous system development, similar to what was observed in the GBM patient analysis (Figure 1d, Figure 2c).

### Ageing hallmarks are present in normal-appearing GBM brain tissue

To further examine the similarity between AD and GBM patient brains at the protein level, we analysed levels of lipofuscin. Accumulation of lipofuscin is a hallmark of ageing and age-related neurodegeneration, including AD^20^. There was a significant (p = 0.0411) increase in lipofuscin in NA-GBM patient brain samples compared to control samples (Figure 3a,b). We also observed a significant increase in phosphorylated Tau (Ser202, Thr205, p-Tau) in NA-GBM patient samples compared to unaffected control individual samples (Figure 3c,d) (p = 0.0152). Tau is a microtubule-associated protein that becomes hyperphosphorylated and forms insoluble aggregates in neurodegenerative tauopathies including AD^21^. To confirm that total p-Tau levels increase in NA-GBM samples we performed western blot analysis and found a significant increase (p = 0.0286) in p-Tau in NA-GBM samples compared to the controls, as well as an expected increase of p-Tau in the AD samples (Figure 3e,f). In contrast, we found no significant increase in Amyloid-β_42_, another well-established hallmark of AD, in NA-GBM samples compared to control samples (Supplementary Figure 4). In summary, beyond the transcriptional similarities these data suggest that the brain of GBM patients contains hallmarks of accelerated ageing and AD

**Figure 3.**
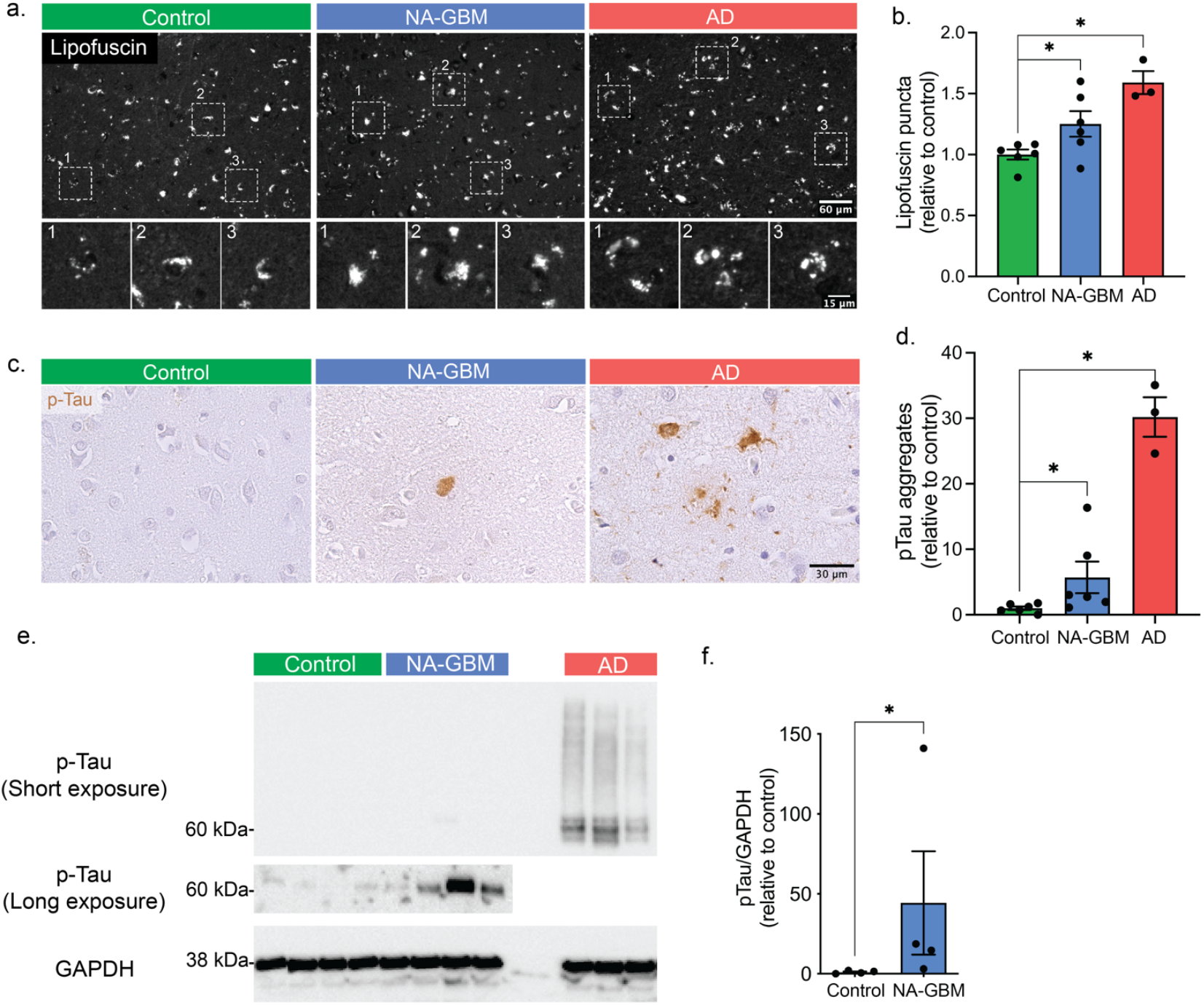
Normal-appearing GBM patient brain tissue shows increased levels of lipofuscin and hyper-phosphorylated tau. a. Immunofluorescence of lipofuscin granules in unaffected control, NA-GBM and AD samples. Top panels scale bar = 60 µm. Bottom panels show boxed area in top panels. Bottom panels scale bar = 15 µm. b. Quantification of number of lipofuscin granules per tile in NA-GBM and AD samples, relative to the control (unaffected control n = 6, NA-GBM n = 6, AD n = 3, NA-GBM vs unaffected control p = 0.0411. AD vs unaffected control p = 0.0238, Mann-Whitney U. * indicates a P value < 0.05). Data are represented as mean ± SEM. c. Immunohistochemistry staining for phosphorylated Tau (Ser202, Thr205) in unaffected control, NA-GBM and AD samples. Scale bar = 30 µm. d. Quantification of the p-Tau IHC staining, showing number of p-Tau aggregates per 9 mm^2^ in NA-GBM and AD samples, relative to control samples (unaffected control n = 6, GBM n = 6, AD n = 3. NA-GBM vs unaffected control p = 0.0152. AD vs unaffected control p = 0.0238, Mann-Whitney U). Data are represented as mean ± SEM. e. Western blot analysis of unaffected control, GBM and AD brain tissue with phosphorylated-Tau (Ser202, Thr205) antibody and GAPDH antibody (low exposure = 5 seconds, high exposure = 355 seconds). f. Western blot quantification showing levels of phosphorylated-Tau relative to GAPDH in unaffected control and NA-GBM brain tissue (p = 0.0286, Mann Whitney U). Data are represented as mean ± SEM.

## Discussion

Our findings reveal that non-tumour regions of GBM patient brains display AD-like ageing hallmarks. One of the factors underlying the AD-like phenotypes observed in GBM patients may be the genotoxic effects of radiotherapy and chemotherapy^22,23^. Indeed, studies have shown an association between DNA damage and a loss of protein homeostasis^24-27^. Remarkably, the proteins that aggregate after genotoxic stress overlap with proteins that aggregate in the background of neurodegenerative disease including AD^27^. DNA damage and reduced expression of DNA damage response proteins have also been implicated in AD^28,29^. However, how DNA damage can trigger a loss of protein homeostasis remains unclear^25,27,28^.

In our study, we observed an increase in p-Tau levels in NA-GBM samples, but absence of amyloid-β aggregates (Figure 3c,d, Supplementary Figure 4). p-Tau forms aggregates in a range of brain pathologies, including AD, progressive supranuclear palsy and corticobasal degeneration^30^. Therefore, an increase in p-Tau suggests several possible root causes of accelerated ageing in the brain of GBM patients, which may differ from AD. An accumulation of p-Tau has previously been shown in a mouse GBM xenograft model^31^. Therefore, we cannot exclude the possibility that some of the ageing phenotypes observed in the surrounding healthy brain tissue might be due to the impact of the tumour itself. A previous meta-analysis study, although focused on the tumour response, also found a transcriptomic overlap between tumour GBM and AD samples^32^. However, in the present study, human brain tissue samples were located far from the tumour site (Supplementary Figure 1a, Table 1). Additionally, cell deconvolution analysis and Nestin staining indicated minimal to no GBM infiltration in our NA-GBM brain samples (Figure 1b-d). Therefore, the direct contribution of GBM cells to the Tau pathology seems unlikely, and indicates that the accelerated ageing and protein aggregation observed in the NA-GBM samples might be a result of the genotoxic effects of chemotherapy and radiotherapy treatments.

Overall, our study demonstrates that the brain of GBM patients display an AD-like accelerated ageing phenotype. Whether this is due to the impact of the tumour itself or a consequence of radiotherapy and/or chemotherapy treatment remains to be further investigated. Furthermore, the results of this work provide the basis for testing therapeutic strategies targeting p-Tau aggregation^33^ in brain tumour patients thereby improving their quality of life.

## Acknowledgments

We thank the UMCG sequencing department for performing the bulk RNA sequencing. We thank W.F.A. den Dunnen from the UMCG pathology department for providing resected tumour tissue from a GBM patient, and for input regarding the amyloid-β staining protocol. This work was funded by KWF Kankerbestrijding (project numbers 12487 and 11148). Post-mortem brain tissues (frozen and fixed) were acquired from the NIH NeuroBiobank (request numbers 1116 and 1834). Schematics in Figure 1 and Supplementary Figure 1 were designed using Biorender.

## Author contributions

L.B. and S.B. designed the study. A.P.A., L.B. and S.B. wrote the paper with input from all authors. A.P.A. performed and analysed the RNA-sequencing experiments. E.G. provided the code and further assistance for RNA-sequencing analysis. B.J.L.E. provided input on gene enrichment analysis and interpretation. A.P.A. performed and analysed the lipofuscin and p-Tau stainings. L.B. and A.P.A. performed and analysed the amyloid-β and Nestin brain stainings. M.K. performed and analysed the immunoblots. L.B. and D.C.V. processed and prepared the human brain tissues for analysis. D.C.V. performed pilot tissue analyses.

## Materials & Methods

### Study design

Brain tissues for this study were obtained from the NIH NeuroBiobank. Post-mortem samples from unaffected control individuals, GBM patients, and AD patients (Braak stages 4-6) were selected based on age (between 44-62 years old) and sex (similar distribution of male and female). Samples were predominantly from the dorsolateral prefrontal cortex whenever possible. Some non-tumour NA-GBM samples were from the cingulate cortex. Unaffected control samples were selected to be accordingly region-matched. Redacted patient medical records were analysed to ensure the GBM patient brain region samples were far from the site of the tumour. The GBM tumour sample for the Nestin staining was obtained from the UMCG pathology department and derived from a 49-year-old female living patient that underwent tumour debulking.

### RNA quality and sequencing

RNA isolation was performed on 12 unaffected control, 15 NA-GBM and 11 AD brain tissue samples. Approximately 40 mg of frozen brain tissue was processed using Qiagen RNA Lipid Tissue kit. Quality of the RNA was determined using TapeStation, only samples with a RIN > 5 were included in the experiment. 70 ng sample RNA was used for library preparation with the Lexogen QuantSeq 3′ mRNA-Seq Library Prep Kit (FWD). cDNA libraries were pooled equimolarly and sequenced on a NextSeq 500 at the sequencing facility in the UMCG.

### Transcriptomic analysis

Data pre-processing was performed with the Lexogen QuantSeq 2.3.1 FWD UMI pipeline on the BlueBee Genomics Platform. The gene count files were imported into R. The ‘boxplot’ function was used to check the read count per million reads distributions in each sample. Samples with a consistent median were selected for further analysis (median = 3 ± 0.25). PCA was performed on normalized read counts and plots were generated in R using ‘ggplot2’. PCA plots were used to check for correlations with library size, age, brain region and PMI. Based on these plots, samples were further narrowed down to those with PMI < 37 hours, and a library size > 800,000. This resulted in a total of five unaffected control, five NA-GBM, and five AD samples for final DEG analysis. DEG analysis was performed using ‘edgeR’^34^. Volcano plots and heatmaps were generated using the CRAN package ‘ggplot2’. Deconvolution analysis was performed using CIBERSORTx^11^, using the dataset from Lake *et al*. 2018^12^ as the signature reference dataset.

### GO analysis and enrichment analysis

GO analysis of DEG heat map clusters was performed using G profiler^35^. GO analysis of all DEGs was performed using WebGestalt (WEB-based GEne SeT AnaLysis Toolkit, RRID:SCR_006786)^36^. Enrichment analysis^37^ of GBM vs unaffected control DEGs was performed by comparing the data set to the “Chemical and Genetic perturbations” data set^38^ using iDEP93^39^, selecting transcriptomic data sets from the top hits for comparison. PGSEA comparisons against the Jensen disease database were also calculated using iDEP93^39,40^. The number of genes that overlap with transcriptomic data of other neurodegenerative diseases was calculated using the ‘match’ function in R. P values were calculated using a hypergeometric distribution test, using the ‘phyper’ function in R.

### Immunofluorescence staining and imaging

Formalin-fixed and paraffin-embedded brain tissue from unaffected control, GBM patient, and AD patient donors were provided by the NIH Biobank. Tissue was cut into 5 µm thick slices and collected on TOMO microscope slides. The staining procedure for lipofuscin was as follows: paraffin-embedded tissue sections were de-paraffinized in xylene and ethanol, then rinsed in demiH_2_O. Slides were washed 3x with PBS, and incubated on DAPI for 10 minutes. Slides were washed again in PBS 3 × 5 minutes, and mounted with Faramount Aqueous Mounting Medium. The autofluorescence of lipofuscin was imaged using a Leica DM6B. Snapshots were made of each sample on a representative grey-matter area of 619.57 µm x 464.68 µm with a 20X magnification. Lipofuscin puncta were automatically quantified using an ImageJ (RRID:SCR_002285) macro:

> “{run(“Enhance Contrast”, “saturated=0.35”);
>
> run(“Apply LUT”);
>
> run(“Auto Threshold”, “method=Otsu white”);
>
> run(“Gaussian Blur…”, “sigma=2”);
>
> setOption(“BlackBackground”, false);
>
> run(“Make Binary”);
>
> run(“Analyze Particles…”, “size=10-Infinity pixel circularity=0.2-1.00 display exclude summarize add”);}”

### Immunohistochemistry

Formalin-fixed and paraffin-embedded brain tissue from unaffected control, GBM patient, and AD patient donors were provided by the NIH Biobank. Tissue was cut into 5 µm thick slices and collected on TOMO microscope slides. The staining procedure for p-Tau was as follows: paraffin-embedded tissue sections were de-paraffinized in xylene and ethanol, then rinsed 1x in demiH_2_O. Antigen retrieval was performed using Histo VT One. Slides were washed 1x in PBS. Peroxidase incubation was performed with 0.5 % H_2_O_2_ in PBS for 30 min at room temperature in the dark. Slides were then washed in PBS 3x 5 minutes. The slides were blocked with blocking buffer (4% rabbit serum, 1 % BSA and 0.1 % Triton) for 1 hour at room temperature. The slides were incubated with the primary antibody diluted in blocking buffer (Phospho-Tau (Ser202, Thr205) Monoclonal Antibody (AT8) cat# MN1020, RRID:AB_223647, used at 1:2000) overnight at 4 ᵒC. Slides were then washed in PBS 3x 5 minutes and incubated for 1 hour with the secondary antibody (biotinylated rabbit anti-mouse, RRID:AB_2687571, used at 1:300) diluted in blocking buffer at room temperature. The slides were incubated an ABC solution (following the Vectastain elite ABC kit, cat# PK-6100) for 30 minutes at room temperature. Slides were then washed in PBS 3x 5 minutes. DAB solution was added under a stereoscope, and time elapsed until visible staining occurred in a positive (AD) sample was timed. The same timing was then used for the other samples. The reaction was stopped with demiH_2_O, and the sections were incubated in hematoxylin, then washed with demiH_2_O for 10 minutes. The sections were dehydrated in an ethanol gradient, mounted with Eukit and dried for 1-2 days.

The staining procedure for amyloid-β was as follows: paraffin-embedded tissue sections were de-paraffinized in xylene and ethanol, then rinsed 1x in demiH_2_O. Antigen retrieval was performed using citric acid and sodium citrate at pH = 6. Slides were washed 3x 5 minutes in PBS, then incubated for 3 min at room temperature with formic acid at room temperature. Slides were washed 3x 5 minutes in PBS. Blocking solution was prepared (1% donkey serum, 1% BSA in PBS). The slides were incubated with the primary antibody diluted in blocking buffer (β-Amyloid Antibody Cell Signalling #2454; rabbit, RRID:AB_2056585, used at 1:500) overnight at 4 ᵒC. Slides were then washed in PBS 3x 5 minutes and incubated for 1 hour with the secondary antibody (biotinylated donkey anti-rabbit, RRID:AB_2340593, used at 1:400) diluted in blocking buffer. The slides were incubated an ABC solution (following the Vectastain elite ABC kit, cat# PK-6100) for 30 minutes at room temperature. Slides were then washed in PBS 3x 5 minutes. DAB solution was added for 3:30 minutes at room temperature. The reaction was stopped with demiH_2_O, and the sections were incubated in hematoxylin, then washed with demiH_2_O for 10 minutes. The sections were dehydrated in an ethanol gradient, mounted with Eukit and dried for 1-2 days.

The staining procedure for Nestin was as follows: paraffin-embedded tissue sections were de-paraffinized in xylene and ethanol, then rinsed 1x in demiH_2_O and 1x in TBST. Antigen retrieval was performed using citric acid and sodium citrate at pH = 6. Slides were washed 3x 5 minutes in PBS, then blocked with 3 % H_2_O_2_ for 10 min at room temperature. Slides were washed 2x 5 minutes in demiH_2_O and 1x 5 minutes in TBST. Slides were blocked with blocking solution (5% goat serum in TBST) for 1 hour at room temperature. Primary antibody incubation was performed overnight at 4 ᵒC (mouse anti-human Nestin cat# MAB 1259, RRID:AB_2251304, used at 1:100). Slides were washed 3x 5 minutes in TBST and then incubated with 3 drops of SignalStain Boost IHC Detection Reagent (HRP, Mouse, Cell Signalling cat# 8125S) for 30 minutes at room temperature. Slides were then washed in TBST 3x 5 minutes. DAB solution was added for 3:30 minutes at room temperature. The reaction was stopped with demiH_2_O, and the sections were incubated in hematoxylin, then washed with demiH_2_O for 10 minutes. The sections were dehydrated in an ethanol and xylene gradient, mounted with Eukit and dried for 1-2 days.

### Quantification of amyloid-β, p-Tau and Nestin

For quantification of amyloid-β, p-Tau and Nestin immunohistochemistry staining, imaging was performed using a Leica DM6B. Tiled snapshots were made of each sample on three representative grey-matter area of approximately 9 mm^2^ with a 20X magnification. Amyloid-β, p-Tau and Nestin aggregates were quantified manually and blindly using the multi-point tool in ImageJ (RRID:SCR_002285).

### Immunoblot analysis of human brain tissue

Frozen human brain tissue was cut into 40 μm thick sections. Next, the sections were lysed in 1x Laemmli buffer. After resuspension, the samples were sonicated and centrifuged at 10.000 rpm for 20 minutes at 4 ᵒC. The supernatant was stored at -80 ᵒC until use. The total protein concentration was determined using a DC Protein Assay Kit (Bio-rad). For protein separation, samples were boiled for 5 minutes and loaded onto TGX FastCast acrylamide gels 10% (Bio-rad).

Proteins were transferred onto nitrocellulose membranes (Bio-rad) and blocked using 10% milk powder in PBST. Next, the membranes incubated overnight at 4 ᵒC with specific antibodies against Phosphorylated Tau (Ser202, Thr205) (mouse, 1:1000, Thermo Fisher, MN1020, RRID:AB_223647), and GAPDH (mouse, 1:10000, Fitzgerald, 10R-G109A, RRID:AB_1285808). Afterwards, the membranes incubated with an anti-mouse HRP-linked secondary antibody (1:5000, GE Healthcare, NXA931, RRID:AB_772209). Either Pierce ECL Western Blotting Substrate (Thermo Fisher) or SuperSignal West Dura Substrate (Thermo Fisher) was used for protein visualization. Images were acquired using a ChemiDoc Imaging System (Bio-Rad) and processed with Image Lab 6.1 software (Bio-Rad).

## Data availability

Bulk RNA-sequencing data are available under GEO number GSE207821. The secure GEO access token is available upon request. Further raw data and analyses are in the supplementary information, or available upon request.

**Supplementary Figure 1.**
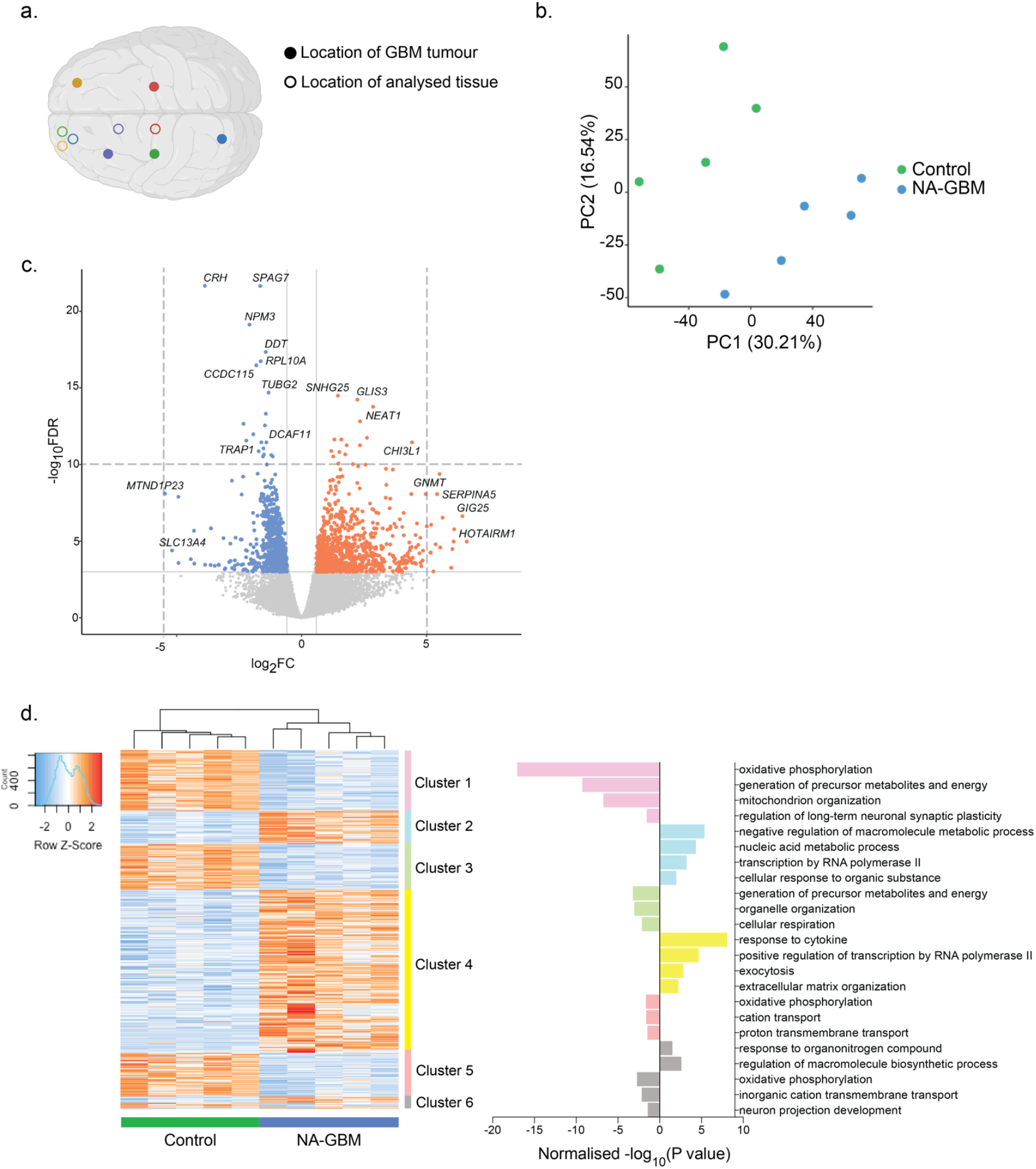
Differential gene expression analysis of NA-GBM and unaffected control samples. a. Schematic of the location of the GBM tumours and the location of the normal-appearing tissue that was analysed (Table 1). Purple = GBM_1, Yellow = GBM_2, Blue = GBM_3, Red = GBM=4, Green = GBM_5. b. The unaffected control (n = 5) and NA-GBM sample (n = 5) RNA-sequencing datasets plotted on a Principal Component Analysis (PCA) plot. c. Volcano plot showing individual DEGs when comparing NA-GBM to unaffected control samples, with fold change > 1.5, and FDR < 0.05. d. Clustered heat map of DEGs when comparing NA-GBM patient samples to unaffected controls (fold change > 1.5 and FDR < 0.05). GO analysis (Biological Processes) of individual clusters using g:Profiler, normalised -log_10_(P value) is plotted on the right.

**Supplementary Figure 2.**
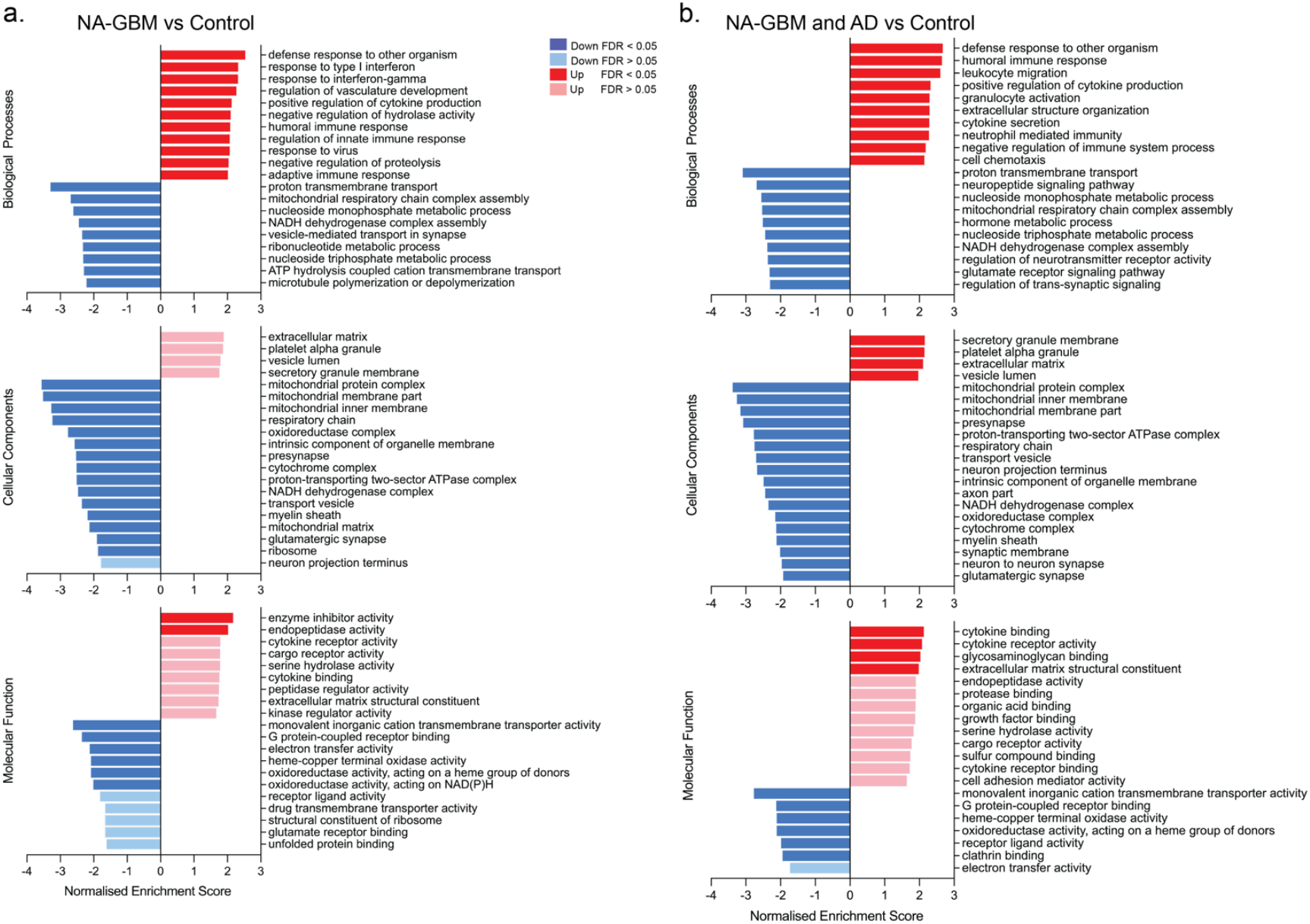
Gene ontology analysis of NA-GBM vs unaffected control DEGs and NA-GBM/AD vs unaffected control DEGs. a. GO analysis of NA-GBM vs unaffected control DEGs, Biological Processes, Molecular Functions and Cellular Components, using WEB-based GEne SeT AnaLysis Toolkit (WebGestalt)^36^. Red = upregulated pathway, blue = downregulated pathway. b. GO analysis of NA-GBM and AD vs unaffected control DEGs, Biological Processes, Molecular Functions and Cellular Components, using WEB-based GEne SeT AnaLysis Toolkit (WebGestalt)^36^. Red = upregulated pathway, blue = downregulated pathway.

**Supplementary Figure 3.**
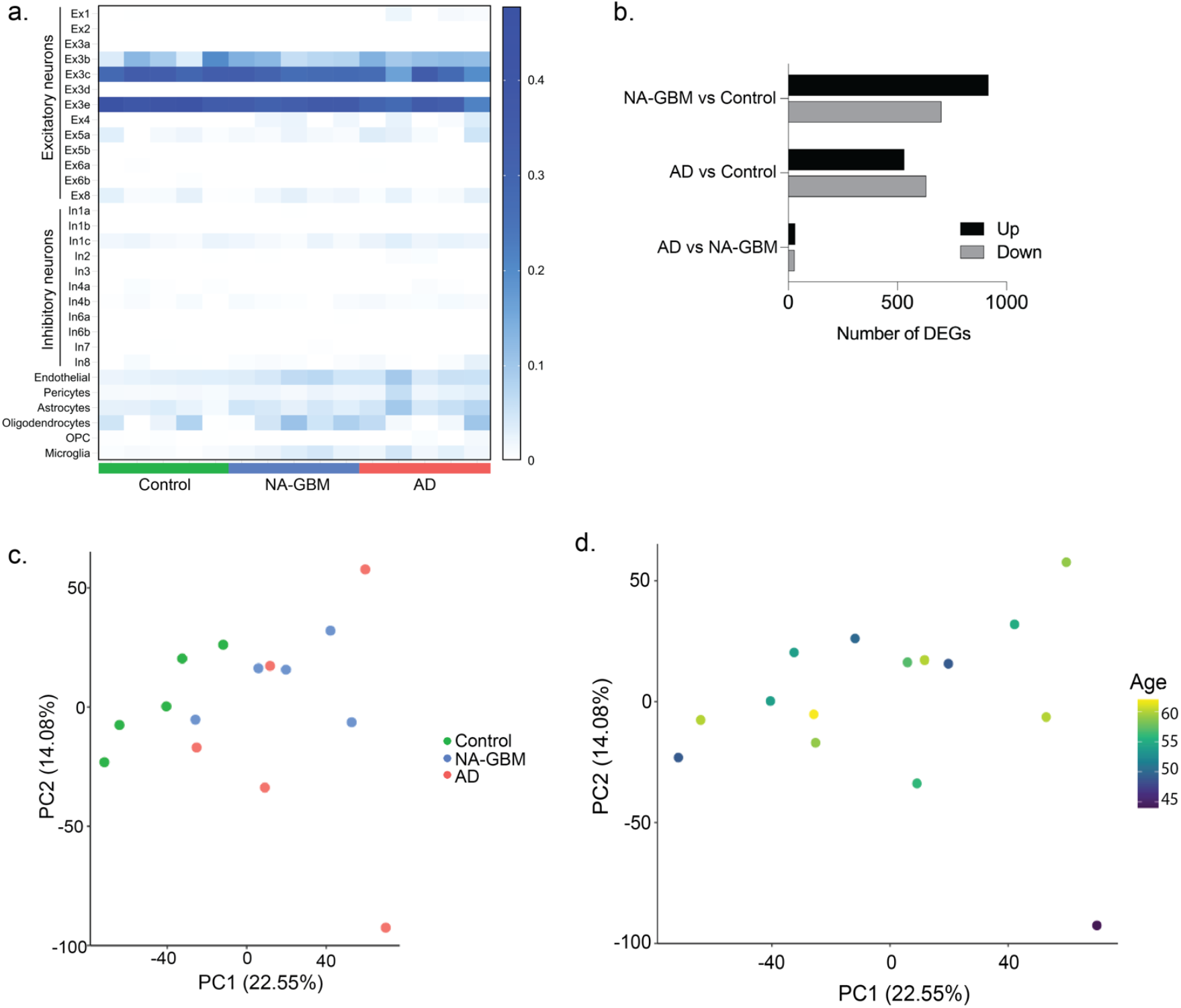
Transcriptomic similarities between NA-GBM and AD patient brain tissue. a. Cell deconvolution analysis of the unaffected control (n = 5), NA-GBM (n = 5) and AD (n = 5) data sets using the CIBERSORTx analytical tool^11^. Heat map indicates inferred ratio of cell types. Endo = endothelial, Peri = pericytes, Astro = astrocytes, Oligo = oligodendrocytes, OPC = oligodendrocyte precursor cells, Micro = microglia. b. Number of DEGs when comparing NA-GBM to control, comparing AD to control, and comparing AD to NA-GBM (fold change > 1.5 and FDR < 0.05). c. Unaffected control (n = 5), NA-GBM (n = 5) and AD samples (n = 5) RNA-sequencing datasets plotted on a PCA plot. d. Unaffected control, NA-GBM and AD samples RNA-sequencing datasets plotted on a PCA plot, colour-coded by age.

**Supplementary Figure 4.**
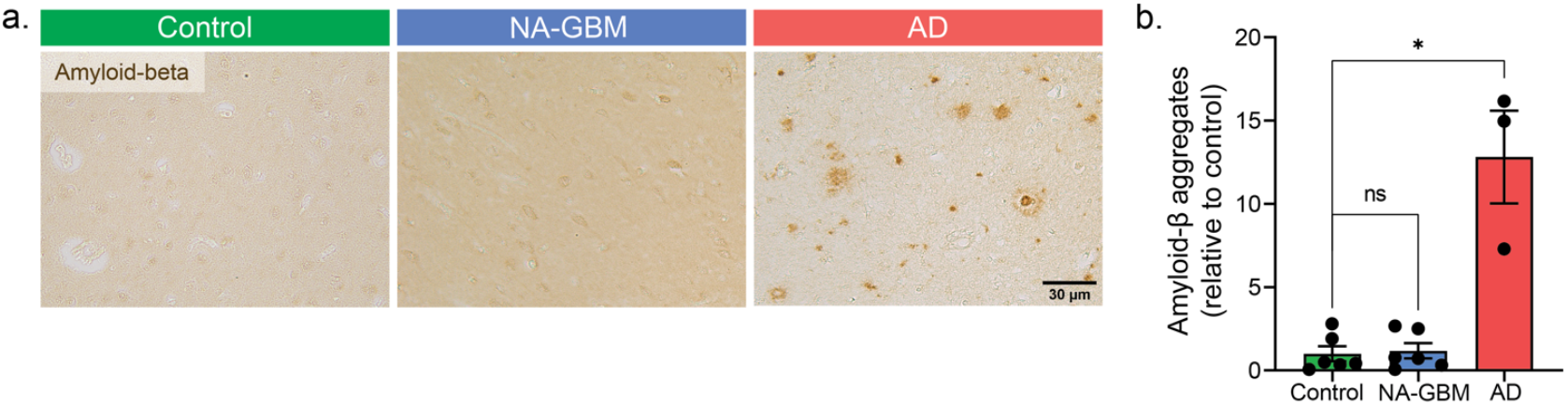
NA-GBM samples do not show any increase in amyloid-β levels. a. Immunohistochemistry staining for amyloid-β in unaffected control, NA-GBM and AD samples. Scale bar = 30 µm. b. Quantification of the amyloid-β staining, showing number of amyloid-β aggregates per 9 mm^2^ in unaffected control, NA-GBM and AD samples, relative to the controls. Unaffected control n = 6, NA-GBM n = 6, AD n = 3 (GBM vs unaffected control p = 0.8182, AD vs unaffected control p = 0.0238, Mann Whitney U). Data are represented as mean ± SEM.

## Supplementary Data 1

List of DEGs in multiple analyses (fold change > 1.5, FDR < 0.05):

a. NA-GBM patient vs unaffected control brain tissue.
b. AD patient vs unaffected control brain tissue.
c. NA-GBM and AD patient vs unaffected control brain tissue.
d. NA-GBM patient vs AD patient brain tissue.
e. Overlapping DEGs between NA-GBM vs unaffected control and AD vs unaffected control.

